# GreenGate 2.0: backwards compatible addons for assembly of complex transcriptional units and their stacking with GreenGate

**DOI:** 10.1101/2023.08.02.551719

**Authors:** Marcel Piepers, Katarina Erbstein, Jazmin Reyes-Hernandez, Changzheng Song, Tomas Tessi, Vesta Petrasiunaite, Naja Faerber, Kathrin Distel, Alexis Maizel

## Abstract

Molecular cloning is a crucial technique in genetic engineering that enables the precise design of synthetic transcriptional units (TUs) and the manipulation of genomes. GreenGate and several other modular molecular cloning systems were developed about ten years ago and are widely used in plant research. All these systems define grammars for assembling transcriptional units from building blocks, cloned as Level 0 modules flanked by four-base pair overhangs and recognition sites for a particular Type IIs endonuclease. Modules are efficiently assembled into Level 1 TUs in a hierarchical assembly process, and Level 2 multigene constructs are assembled by stacking Level 1 TUs. GreenGate is highly popular but has three main limitations. First, using ad-hoc overhangs added by PCR and classical restriction/ligation prevents the efficient use of a one-pot, one-step reaction to generate entry clones and domesticate internal sites; second, a Level 1 TU is assembled from a maximum of six modules, which may be limiting for applications such as multiplex genome editing; third, the generation of Level 2 assemblies is sequential and inefficient. GreenGate 2.0 (GG2.0) expands GreenGate features. It introduces additional overhangs, allowing for the combination of up to 12 Level 0 modules in a Level 1 TU. It includes a Universal Entry Generator plasmid (pUEG) to streamline the generation of Level 0 modules. GG2.0 introduces GreenBraid, a convenient method for stacking transcriptional units iteratively for multigene assemblies. Importantly, GG2.0 is backwards compatible with most existing GreenGate modules. Additionally, GG2.0 includes Level 0 modules for multiplex expression of guide RNAs for CRISPR/Cas9 genome editing and pre-assembled Level 1 vectors for dexamethasone-inducible gene expression and ubiquitous expression of plasma membrane and nuclear fluorescent markers. GG2.0 streamlines and increases the versatility of assembling complex transcriptional units and their combination.

## Introduction

Molecular cloning is an integral part of genetic engineering. It allows for the precise design of synthetic transcriptional units (TUs), the manipulation of genomes and the alteration of gene expression. Several methods allow the modular combination of pre-generated modules into destination vectors and were developed in the last 20 years [1]. The Gateway system, which relies on proprietary BP/LR recombinases, allows in its latest iteration to combine up to four DNA fragments in a destination plasmid [2]. The Gibson assembly [3], or the conceptually similar In-fusion system [4], allow the combination of linear DNA fragments sharing stretches of 15–30 base pairs in a single tube reaction combining the use of an exonuclease, polymerase and ligase. Both methods are modular but rely on engineering overlapping sequence fragments between modules. Thus, they do not allow for the re-utilization of DNA modules in multiple and diverse DNA assemblies.

Golden Gate cloning provides the highest level of modularity and reusability of modules [5]. It is based on Type IIs restriction enzymes, which can recognise non-palindromic sequence motifs and cut outside their recognition site, creating defined overhangs on either strand of DNA. This system allows for directional assembly of multiple DNA parts using only one restriction enzyme. The parts are released from their original plasmids and assembled into a new plasmid backbone in a one-pot, one-step reaction without time-consuming steps such as custom primer design, PCR amplification, and gel purification. Modules must be housed between a convergent pair of Type IIs recognition sequences to perform this reaction. The accepting plasmid must have a divergent pair of recognition sequences for the same enzyme. Additionally, the overhangs created by the digestion should be unique and non-palindromic. Finally, the plasmid backbones housing the parts should have a different antibiotic resistance than the plasmid into which parts will be assembled [1].

Over the years, multiple variants of Golden Gate-based cloning toolkits have been developed for plant systems: GreenGate [6], GoldenBraid [7], and Modular Cloning (MoClo) [8,9] were concomitantly developed about ten years ago. Additional systems such as Mobius, Joint Modular Cloning (JMC) and LoopAssembly were recently introduced [10–13]. All these systems define grammars for assembling TUs from building blocks, including promoters, untranslated regions, signal peptides, coding sequences, and terminators. These building blocks are cloned as Level 0 modules flanked by the four-base pair overhangs and recognition sites for a particular Type IIs endonuclease. In a second hierarchical level (Level 1), modules are efficiently assembled into TUs using Golden Gate cloning. Multigene constructs are assembled with another Golden Gate reaction at a higher hierarchical level (Level 2). A prerequisite for any Golden Gate-based DNA assembly is removing internal recognition sites from the brick for the relevant Type IIs endonucleases, a process called “sequence domestication”. All systems feature sets of modules for commonly used plant promoters, transcriptional terminators, epitope tags and reporter genes [6,9,14] available via AddGene, allowing end users to jump into hierarchical DNA assemblies quickly.

The different systems have their specificities. While GreenGate relies on a single Type IIs enzyme (BsaI, [6]) for the hierarchical assembly of constructs, MoClo, Loop, JMC, and Mobius cloning systems rely on the iterative use of two different Type IIs enzymes and GoldenBraid and JMC on three. MoClo uses BsaI and BpiI [9], Mobius uses BsaI and AarI [10], Loop BsaI and SapI [13] and GoldenBraid BsaI, BsmBI and BtgZI [14], JMC uses AarI/PaqCI, BsaI and Esp3I [12]. Although increasing the requirement for sequence modifications during domestication, this streamlines cloning procedures and increases flexibility. MoClo, GoldenBraid, JMC, Loop, and Mobius designers have agreed on a standard molecular syntax to promote the sharing and re-utilisation of DNA modules for bioengineering purposes, rendering these systems largely, though not entirely, compatible [1]. The different systems differ in how many Level 0 modules can be combined in a Level 1 assembly. GreenGate and MoClo allow the combination of up to six modules, while GoldenBraid, JMC, Loop, and Mobius up to 11. The creation of Level 0 modules can rely on a set of entry vectors containing the Type IIs overhangs specific for a given module (promoter, coding sequence, terminator), as is the case for GreenGate and MoClo. GoldenBraid and Mobius use a polyvalent entry vector system where PCR incorporates the four-base pair overhangs into the modules. The systems additionally differ in the way they assemble multigene constructs. Assemblies can be linear (GreenGate, MoClo and JMC) or cyclical (GoldenBraid, Loop, Mobius). For linear assemblies of two TUs, GreenGate relies on cloning the TUs in intermediary Level 1 vectors retaining Eco31I sites, which are combined in the final destination Level 2 plasmid. GreenGate can also linearly assemble TUs using a special adapter module and exploiting the sensitivity of the BsaI sites to methylation [6]; Both processes are, however, inefficient relative to a standard GreenGate reaction. MoClo and JMC allow the assembly of up to five (MoClo) and seven (JMC) Level 1 modules in a single step [9,12]. Maximum flexibility is achieved by GoldenBraid, Loop and Mobius, which uses an iterative combinatorial process for multigene construct assembly. GoldenBraid2.0 and Loop use a set of four plasmids (eight for Mobius) to iteratively assemble multigene constructs, either two-by-two (for GoldenBraid) or four-by-four (for Mobius and Loop) [10,13,14].

Here, we introduce GreenGate 2.0 (GG2.0), a backwards-compatible add-on to the GreenGate system that expands the current GreenGate features. GG2.0 introduces an expanded repertoire of overhangs which allow the combination of up to 12 Level 0 modules in a Level 1 TU. GG2.0 introduces a Universal Entry Generator plasmid (pUEG) that streamlines the generation of Level 0 modules. GG2.0 introduces GreenBraid, a convenient method to iteratively stack TUs for multigene assemblies. GG2.0 is compatible with most readily existing GreenGate Level 0 modules. GG2.0 also features a set of Level 0 modules for multiplex expression of guide RNAs for CRISPR/Cas9 genome editing and pre-assembled Level 1 vectors for ubiquitous expression of plasma membrane and nuclear fluorescent markers as well as dexamethasone-inducible gene expression. The GG2.0 plasmids are available as a kit (Addgene XXXX^1^).

## Material and methods

### Preparation and Transformation of Bacterial Strains

For the growth of E. coli cells (DH5α NEB or TOP10), 4 ml of LB growth medium supplemented with appropriate antibiotics were used for overnight incubation at 37°C with constant agitation at 230 rpm. To generate home-made ultra-competent cells of DH5α (NEB) or TOP10 (Thermo Scientific), the Inoue Method [15] was employed. The competent cells were then transformed with the constructs as follows: 1-5 μl of the DNA solution was mixed with 50 μl of the competent cells on ice for 30 minutes, followed by a heat shock at 42°C for 30 seconds, and then re-cooled on ice for 2 minutes. Subsequently, LB medium (250-500 μl) was added, and after 1 hour of incubation at 37°C with constant agitation at 230 rpm, 50 μl of the cell suspension was plated on LB agar plates with antibiotic selection. The plates were then incubated overnight at 37°C. For the transformation of Agrobacterium tumefaciens (GV3101), a heat shock method was used. Specifically, 5 μl of DNA solution was combined with 50 μl of competent cells on ice for 10 minutes, followed by freezing in liquid nitrogen for 5 minutes. A heat shock was then applied by incubating for 5 minutes at 37°C. Afterward, LB medium (800 μl) was added, and the cells were incubated for 3-4 hours at 28°C with constant agitation at 230 rpm. Finally, cells (500-700 μl) were plated on LB agar plates with antibiotic selection.

### Molecular biology reagents and methods

Restriction enzymes were obtained from New England Biolabs (NEB) and Thermo Scientific/Fermentas. T4 Ligase and T4 polynucleotide kinase was obtained from NEB. Plasmids were isolated using the alkaline lysis method [15]. PCRs were performed with the Q5 High-Fidelity Polymerase (NEB). PCR products and DNA fragments containing DNA were cleaned up with the GeneJET PCR Purification Kit (Thermo Scientific) or Nucleospin gel and PCR cleanup kit.

### Part and vector generation

pUEG was obtained by gene synthesis. For the GreenBraid vectors, we replaced the cloning sites of the pCAMBIA-based GoldenBraid vectors ([10,13,14]) with GreenBraid cloning sites by restriction-ligation of annealed and T4-phosphorylated oligos in EcoR1-opened pDGB3α1 for pGB-Zs and BamH1-opened pDGB3Ω1 for pGB-Ys. Ligations were transformed into *E*.*coli* (DH5α), and correct clones were identified through plasmid extraction, digestion and sequencing. The chromogenic bacterial selection markers were added to each GreenBraid vector (spisPink for Z1, Z2, Z1r, and Z2r; sfGFP for Y1, Y2, Y1r, and Y2r). The chromogenic marker genes (sfGFP & spisPink, ([10–13])) were amplified by PCR and inserted in the GreenBraid vectors by restriction-ligation between PstI and BamHI sites. The GreenBraid Z and Y jokers (Z1j, Z2j, Y1j, and Y2j) were obtained by NcoI and XhoI restriction-ligation of 200-208 bp joker sequences that were PCR amplified from GoldenBraid dummy vectors (pDGB1α1_SF for Z1j, pDGB1α2_SF for Z2j, pDGB1Ω1_SF for Y1j, and pDGB1Ω2_SF for Y2j). The pre-assembled Level 1 GreenBraid vectors containing plasma membrane markers, nucleus markers, or inducible expression were constructed following the Level 1 assembly protocol. The Level 0 entry clones containing a dummy 20 bp sequence were obtained by ligating annealed oligonucleotides in an EcoRI + HindIII digested pGGA-000 GreenGate entry clone [6]. The exactitude of the insertion was verified by sequencing.

### Generation of Level 0 entry clones

A detailed protocol for the domestication of endogenous sites and generation of Level 0 entry clones using the Universal Entry Generator (pUEG) can be found in the supplemental material.

### Generation of entry vectors for gRNA expression

Two sets of entry vectors for gRNA expression were generated: one using the U6 promotor (pGG(C1/ C3/ D1/ D3/ E2)-002); the second one uses the U3 promoter (pGG(C2/ C4/ D2A/ E1)-002). In both sets, the promoter is followed by two AarI sites, the scaffold and a terminator. For the pU6 set, pU6, AarI sites, scaffold and the terminator were amplified from a synthetic template (22AACPBF) with primers specific for the respective Level 0 entry vectors (Supplementary Table S2). The respective Level 0 entry vectors and amplified inserts were combined by restriction-ligation with Eco31I and then transformed into *E. coli*, tested by colony PCR and verified by sequencing. For pU3-containing entry vectors, pU3 and the scaffold+term were independently amplified by PCR from pRU42 [16] to add the two AarI sites and then combined by overlap-extension PCR to generate the pU3-AarI-AarI-scaffold-term. These inserts were cloned by Eco31I restriction-ligation in pGGC2, pGGC4, pGGD2A and pGGE1 entry vectors and then transformed into *E. coli*, tested by colony PCR and verified by sequencing.

### GreenGate assembly

A detailed protocol for assembling Level 1 constructs from Level 0 entry clones can be found in the supplemental material. Briefly, the assembly is performed with a one-tube reaction with a total volume of 20 μl, with 1.5 μL of each entry vector (pGGA, pGGB, pGGC, pGGD, pGGE, pGGF all at ∼100 ng/μL), 1 μL of destination vector (∼100ng/μL), 2 μL ATP (NEB, B0202S), 2 μL 10X restriction digest buffer (Thermo Scientific, BY5), 0.5 μL T4 DNA ligase (30U/μL Thermo Scientific, EL0013), 0.5 μL FastDigest Eco31I (Thermo Scientific, FD0294) and 5 μL water. Destination vectors were either pGGZ003 (for original GreenGate cloning) or pGBZ1 (pMP116) or pGBZ2 (pMP117) for GreenGate 2.0. The following thermocycler program was used for the GreenGate assembly: [(37 °C - 05:00; 16 °C - 05:00) x 50; 50 °C - 05:00; 80 °C - 05:00; 4 °C - ∞]. The reaction was performed overnight, and in the next morning, 0.5 μL FastDigest Eco31I was added to the reaction and incubated for one hour at 37 °C, followed by a 15 min-long heat-inactivation step at 80 °C.

### GreenBraid assembly

A detailed protocol for the stacking of TUs, using the GreenBraid (pGB) Level 2 and Level 1 destination vectors, can be found in the supplemental material. Briefly, the assembly is performed with a one-tube reaction with a total volume of 20 μl, with 100 ng of each Level 1 assembly in pGBZ1 (pMP116) and pGBZ2 (pMP117), 50 ng of destination vector pGBY1 (pMP118) or pGBY2 (pMP119), 2 μL of 10x T4 DNA Ligase Buffer with 10 mM ATP (NEB, B0202S), 1 μL of PaqCI/AarI activator (5 pmol, NEB, S0532S), 1 μL of 10 U PaqCI/AarI (NEB, R0745S) and 1 μL of 400 U T4 DNA Ligase (NEB, M0202S). The following thermocycler program was used: [(37 °C - 01:00; 16 °C - 01:00) x 60; 37 °C - 05:00; 60 °C - 05:00; 4 °C - ∞]. Post-assembly, 2 μL of rCutSmart Buffer (NEB, B6004S), 1 μL of PaqCI/AarI activator (5 pmol) and 0.5 μL of PaqCI/AarI were added again and incubated for 1 h at 37 °C. PaqCI/AarI was heat-inactivated by incubating the assembly mix for 20 min at 65 °C.

### Plant material, growth conditions

Seeds of Arabidopsis thaliana ecotype Colombia (Col-0) were surface sterilised (ethanol 70% and SDS 0.1%) and placed on ½ Murashige and Skoog (MS) medium adjusted to pH 5.7 containing 1% agar (Duchefa). Following stratification (4°C in the dark, > 24 h), plants were grown under fluorescent illumination (90 μE m-2 s-1) in long-day conditions (16 h light / 8 h dark) at 22 °C.

### Plant transformation

*Agrobacterium tumefaciens-*based plant transformation was done using the floral dip method [17].

### Microscopy

Confocal Laser-Scanning Microscopy (CLSM) was performed on a Leica SP5 confocal microscope with a 63x, NA = 1.2 water immersion objective. mTurquoise2 fluorescence was detected using the 458 nm excitation laser line and a detection range of 470-510 nm. mVenus fluorescence was detected using the 514 nm excitation laser line and a detection range of 520-540 nm. mScarlet fluorescence was detected using the 561 nm excitation laser line and a detection range of 570-650 nm.

## Results and Discussion

### Overview of GreenGate 2.0

The modular cloning system GreenGate uses Eco31I (BsaI) to combine six individual Level 0 entry vectors into a destination vector (Level 1) in a single reaction. These Level 1 constructs can subsequently be used for the expression in plants. Over the years, hundreds of individual entry vectors have been created by many laboratories which can easily be exchanged and combined. Two TUs can be combined into a single destination vector (Level 2) through an adaptor module, and more than two TUs can be iteratively assembled. In its first iteration, GreenGate suffers from three main limitations. First, to generate a Level 0 part, the ad-hoc overhangs are added by PCR and the part is cloned in a dedicated entry clone for the specific part by classical restriction/ligation using Eco31I. This process prevents using the efficient Golden Gate one-pot, one-step reaction for generating the Level 0 clone and the easy domestication of internal sites [1]. Second, a GreenGate Level 1 transcriptional unit typically consists of six basic modules: Promoter (A), N-tag (B), Coding (C), C-tag (D), Terminator (E) and a Selection marker (F). While this covers most of the typical needs, it is limiting if, for example, more individual modules must be combined to express several guide RNAs for the multiplex editing of several genes [18]. Third, the generation of Level 2 assemblies, either through combining two TUs pre-assembled in intermediary vectors or the iterative addition of one TU, relies on using Eco31I, making the process inefficient [6].

GG2.0 addresses these limitations. It introduces a Universal Entry Generator plasmid that relies on a second Type IIs enzyme (PaqCI / AarI) to enable a one-pot, one-step assembly of several elements to domesticate internal sites and create Level 0 modules easily; GG2.0 expands the repertoire of available overhangs in its grammar to enable the assembly of Level 1 vectors from up to 12 Level 0 modules; Finally, GG2.0 features a new set of destination vectors to use as Level 1 expression vectors in plants. These new destination vectors, owing to the use of a second Type IIs enzyme (PaqCI / AarI), also enable the iterative combination of Level 1 TUs into Level 2 multi-gene constructs with limitless TU stacking potential (a process dubbed GreenBraiding). Importantly, GG2.0 is backwards compatible with existing GreenGate modules and, in most cases, does not require re-cloning existing modules.

### A universal Level 0 entry vector

GG2.0 features a Universal Entry Generator plasmid (pUEG) designed to serve as a polyvalent entry vector for all the GreenGate modules regardless of their category (Figure 2). The desired part is PCR-amplified as a single fragment or, if restriction sites need to be domesticated, as multiple fragments (Figure 2). pUEG contains a pair of divergent PaqCI/AarI sites flanked by overlapping convergent Eco31I recognition sequences. We chose PaqCI/AarI due to its rare cutting ability, specifically recognising the 7bp sequence CACCTGC(4/8)^ and generating a 4bp overhang. This characteristic reduces the necessity for additional modifications during the assembly process while ensuring efficient assembly. The oligonucleotide primers used for amplification add 5’ PaqCI recognition sites and the GreenGate overhang of the desired module. The amplified part(s) and pUEG are combined to generate a GreenGate entry clone in a one-pot GoldenGate digestion–ligation reaction using PaqCI. Domestication of an internal site(s) or combination of several fragments is possible by combining PCR fragments flanked by overlapping PaqCI sites (Figure 2). pUEG contains the chromoprotein amilCP [13] as a visible negative cloning screening marker, which imbues a purple colour after overnight incubation to the colonies that do not contain the insert. The penetrance of the stain was strain-dependent. Indeed, when pUEG transformed DH5α or TOP10 bacteria, all DH5α colonies presented a strong deep purple colouration, while some white colonies could be observed in TOP10 after overnight incubation. Screening with chromogenic proteins presents several advantages. It eliminates the use of IPTG and X-gal, two expensive chemicals. Furthermore, it allows the use of *E. coli* strains that do not contain the lacZΔM15 deletion mutation necessary for X-Gal screening.

**Figure 1.**
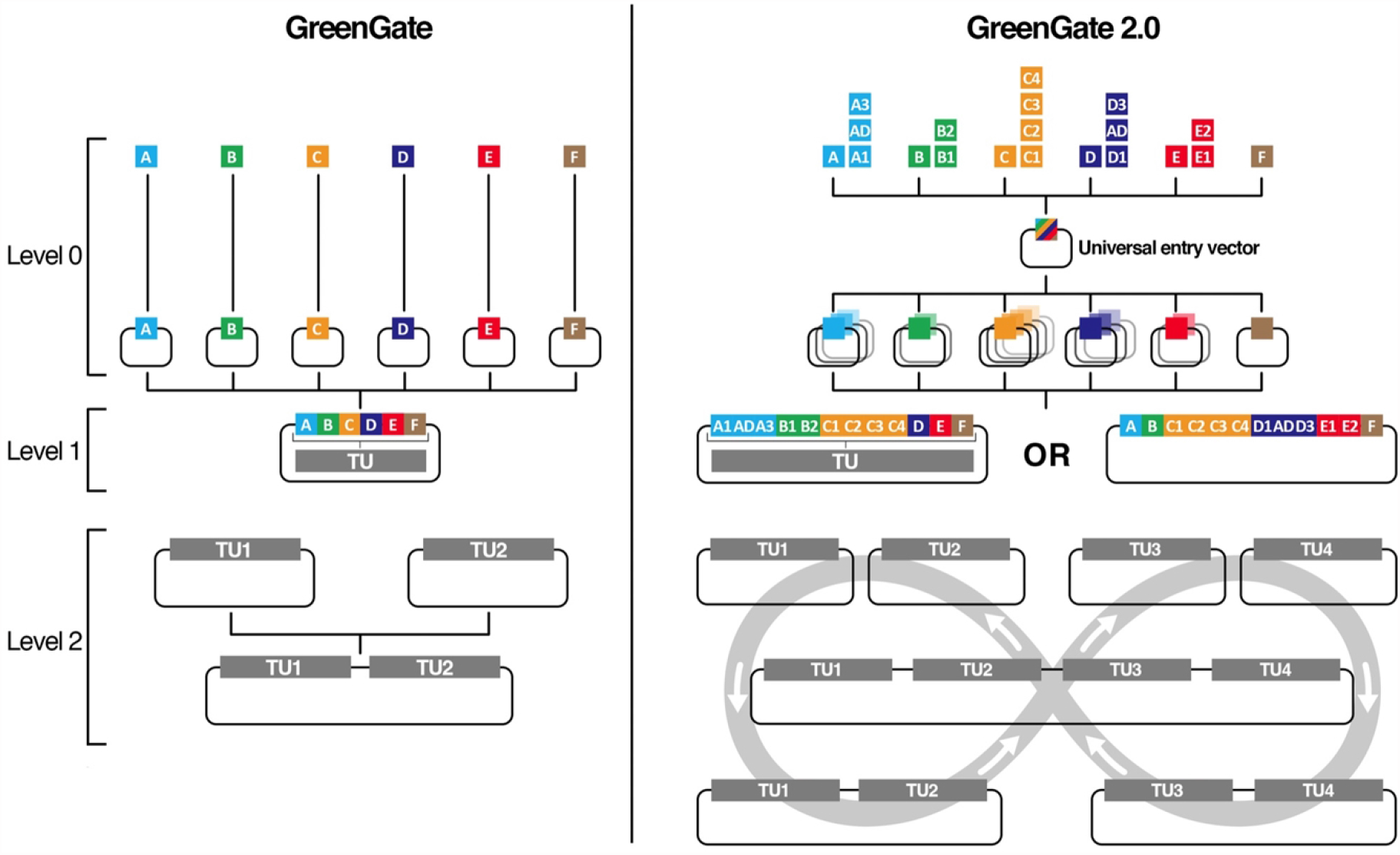
Comparison of GreenGate 2.0 to GreenGate. GreenGate 2.0 (GG2.0) expands the current GreenGate (GG)features set. GG2.0 offers an expanded repertoire of overhangs subdividing the existing GG A to E modules into 14 sub-modules. These modules can be cloned in a unique Universal entry vector to generate Level 0 entry clones. Up to 12 Level 0 clones can be assembled in one GG2.0 transcriptional unit compared to six for GG. Whereas GG only allowed the combination of two TUs to form a multi-gene Level 2 assembly, GG2.0 leverages GreenBraiding for the parallel, iterative assembly of a limitless number of TUs in Level 2 assemblies.

**Figure 2.**
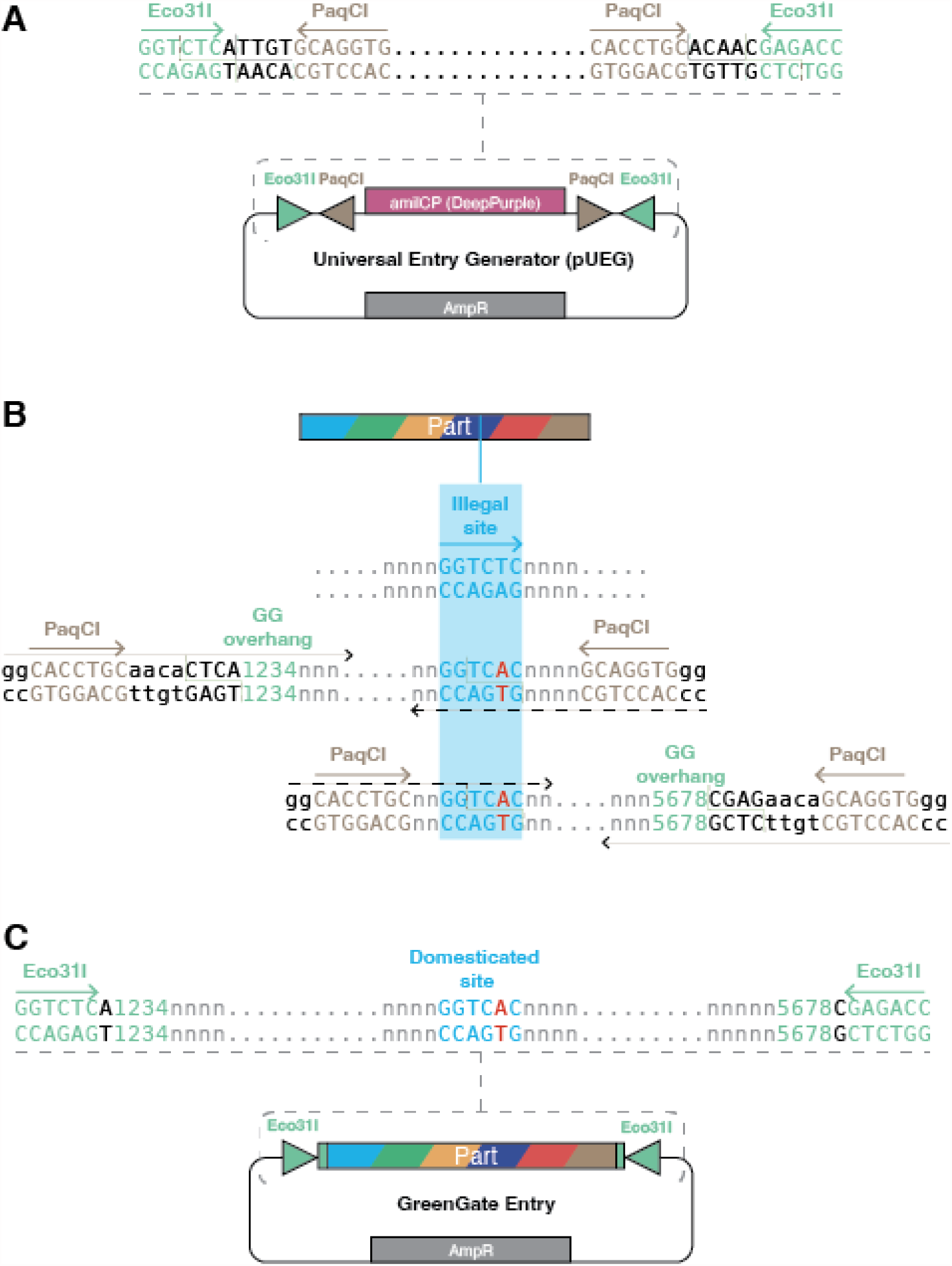
A universal Entry Generator. (A) The Universal Entry Generator plasmid (pUEG) comprises a small backbone conferring resistance to ampicillin in bacteria. It contains a cloning site consisting of a pair of divergent PaqCI/AarI sites flanked by overlapping convergent Eco31I recognition sequences. The plasmid carries a bacterial expression cassette for the chromoprotein AmilCP that renders deep-purple colonies. (B) A sequence containing an illegal Eco31I recognition sequence can be amplified in two fragments using oligonucleotide primers with overhangs (black dashed lines) that (i) introduce a mutation to destroy the illegal site, (ii) add PaqCI/AarI recognition and fusion sites to allow one-step digestion–ligation into the Universal Entry Generator plasmid, and (iii) add the desired GreenGate overhangs (green numbers) that will define the type of GreenGate entry clone. (C) When the resulting amplicons are cloned into pUEG with one-step digestion–ligation, the new standard part will be flanked by a pair of convergent Eco31I recognition sequences capable of releasing the part with the desired GreenGate overhangs (green).

### An expanded repertoire of Level 0 modules

To provide higher granularity and modularity during the assembly, we added six new overhangs to the GreenGate repertoire, which allowed the subdivision of existing GreenGate A, B, C, D and E modules into 14 submodules (Figure 3). We created 21 Level 0 entry vectors that use the same backbone as GreenGate entry clones and contain two convergent Eco31I sites that flank a unique 20 bp sequence and can thus be used, if necessary, as dummy modules during a Level 1 assembly. As several new entry clones share the same new overhangs, not all GG2.0 Level 0 modules can be freely combined in a Level 1 assembly. The GreenGate2.0 modules A1-A3, B1-B2, C1-C4 (ABC set) can be combined with the GreenGate modules D, E and F; alternatively, the GreenGate modules A, B and F can be assembled with the GG2.0 modules C1-C4, D1-D3 and E1-E2 (CDE set). This new set allows up to 12 Level 0 modules to be assembled in a Level 1 TU, dramatically increasing the modularity of GreenGate assemblies. Entry clones spanning multiple overhangs are also available to reduce the complexity of the assembly when a particular module is not required. As up to 12 Level 0 modules and one destination are used in Level 1 TU assemblies, we optimised the protocol and used a higher concentration of T4 ligase. The efficacy of Level 1 assembly remained high when eight or more Level 0 clones were assembled (Table 1). Although we noticed a trend towards a lower efficiency when ten or more entry vectors were combined, the proportion of positive clones remained above 50%.

**Table 1.** Efficacy of Level 1 assembly. N denotes the number of independent assemblies

| Number of Level 0 clones in assembly | Proportion of correct colonies (mean ± SD, N) |
| --- | --- |
| 8 | 72±10%, N=3 |
| 9 | 77±32%, N=2 |
| 10 | 57±12%, N=5 |
| 11 | 47±21%, N=3 |
| 12 | 56±30%, N=4 |

**Figure 3.**
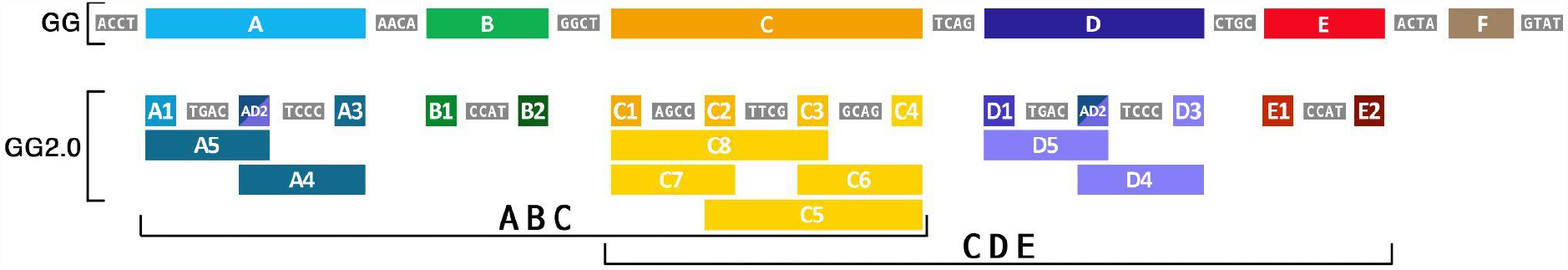
Expanded set of Level 0 modules. GG2.0 introduces six new overhangs (in grey) that subdivide the existing GG A to E modules into 14 submodules (coloured boxes). Bridging modules are provided for increased modularity. As the new overhangs are shared between the submodules A/D and B/E, the GG2.0 modules A1-A3, B1-B2 and C1-C4 (ABC set) can be assembled with the GG modules D, E and F. The GG modules A, B and F can be assembled with the GG2.0 modules C1-C4, D1-D3 and E1-E2 (CDE set) to form Level 1 transcriptional units from up to 12 Level 0 modules.

### A set of Level 0 entry plasmids for multiplex CRISPR/Cas9 genome editing

Taking advantage of the extended set of Level 0 modules, we designed nine Level 0 entry plasmids to express gRNAs from Pol III promoters. These plasmids are in the CDE set and contain the U3 or U6 Pol III promoter, two PaqCI/AarI sites, a Cas9 scaffold sequence and a Pol III terminator (Supplementary Table S1). One or two gRNAs can be cloned by a GoldenGate reaction using PaqCI/AarI. For single gRNAs, annealed oligonucleotides containing the gRNA sequence flanked by the ATTG and GTTT overhangs and the PaqCI/AarI sites are used. For two gRNAs, an amplicon containing the two gRNAs, the scaffold sequence, a Pol III terminator and the pU6 Pol III promoter flanked by PaqCI/AarI is used (Figure 4). The PaqCI/AarI overhangs are common to all entry clones for maximum modularity, so the user can insert the gRNAs in any entry clone. Once loaded with a gRNA, these entry clones can be combined in a single GreenGate assembly to generate a Level 1 T-DNA vector containing up to 18 gRNAs and a promoter-Cas9-terminator cassette for multiplex CRISPR/Cas9 genome editing.

**Figure 4.**
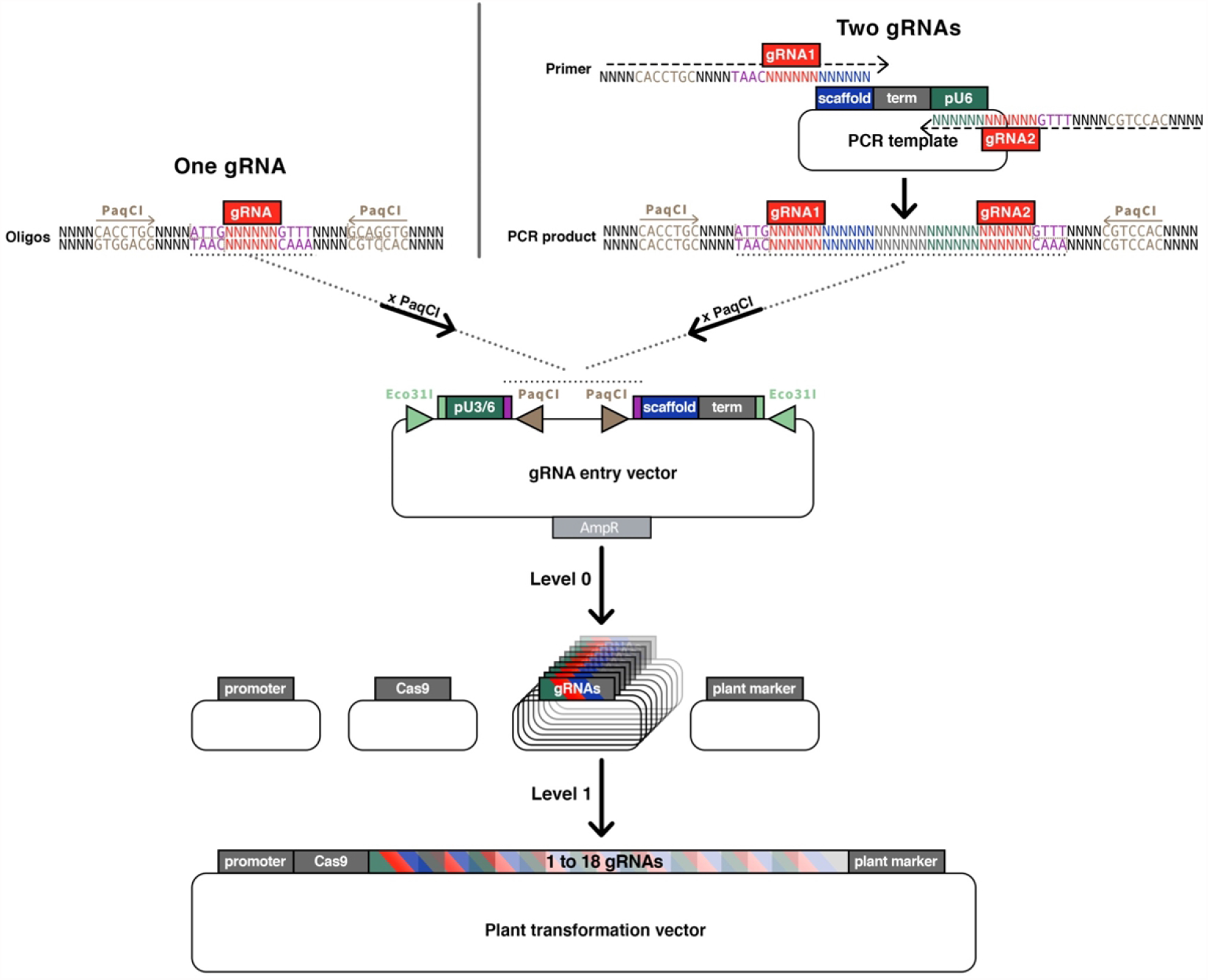
A set of Level 0 entry plasmids for multiplex CRISPR/Cas9 genome editing. A set of GGv2 entry vectors allows the expression of one or two gRNAs from the pU3 or pU6 promoter. Once digested with PaqCI, the vector can receive either a single gRNAs directly ligated as annealed oligonucleotides or two gRNAs if generated by PCR.

### Iterative stacking of TUs using GreenBraid vectors

The original GreenGate cloning offers two methods for stacking TUs on a single T-DNA. The first involved assembling individual TUs in intermediate vectors combined in a second assembly reaction in a regular destination Level 2 vector. The necessity to maintain the Eco31I sites in the intermediate vectors led to reduced assembly efficiency for single TUs compared to a regular GreenGate assembly. Additionally, the intermediate vectors were unsuitable for Agrobacterium-mediated plant transformation. The second method enabled the assembly of more than two TUs on a single T-DNA thanks to a particular F adapter module that exploited the sensitivity of Eco31I sites to cytosine methylation to sequentially add one TU to an existing Level 1 vector. This sequential method was increasingly time-consuming as the number of desired TUs increased and was associated with fewer transformants or a higher number of false-positive colonies than a regular GreenGate assembly [6].

To circumvent these limitations, GG2.0 introduces a set of plant-compatible Level 1 and Level 2 destination vectors that enable the parallel and iterative two-by-two stacking of TUs, a process called GreenBraiding (Figure 5). GreenBraiding is a two-step process that can be looped indefinitely. The assembly of single TUs takes place in Level 1 vectors that can be combined to form a Level 2 vector with two TUs. Two Level 2 vectors can be assembled to combine four TUs and generate back a Level 1 destination vector. By alternating between Level 1 and Level 2 vectors (Figure 5B), GreenBraiding enables the stacking of any number of TUs as far as the vector can handle.

**Figure 5.**
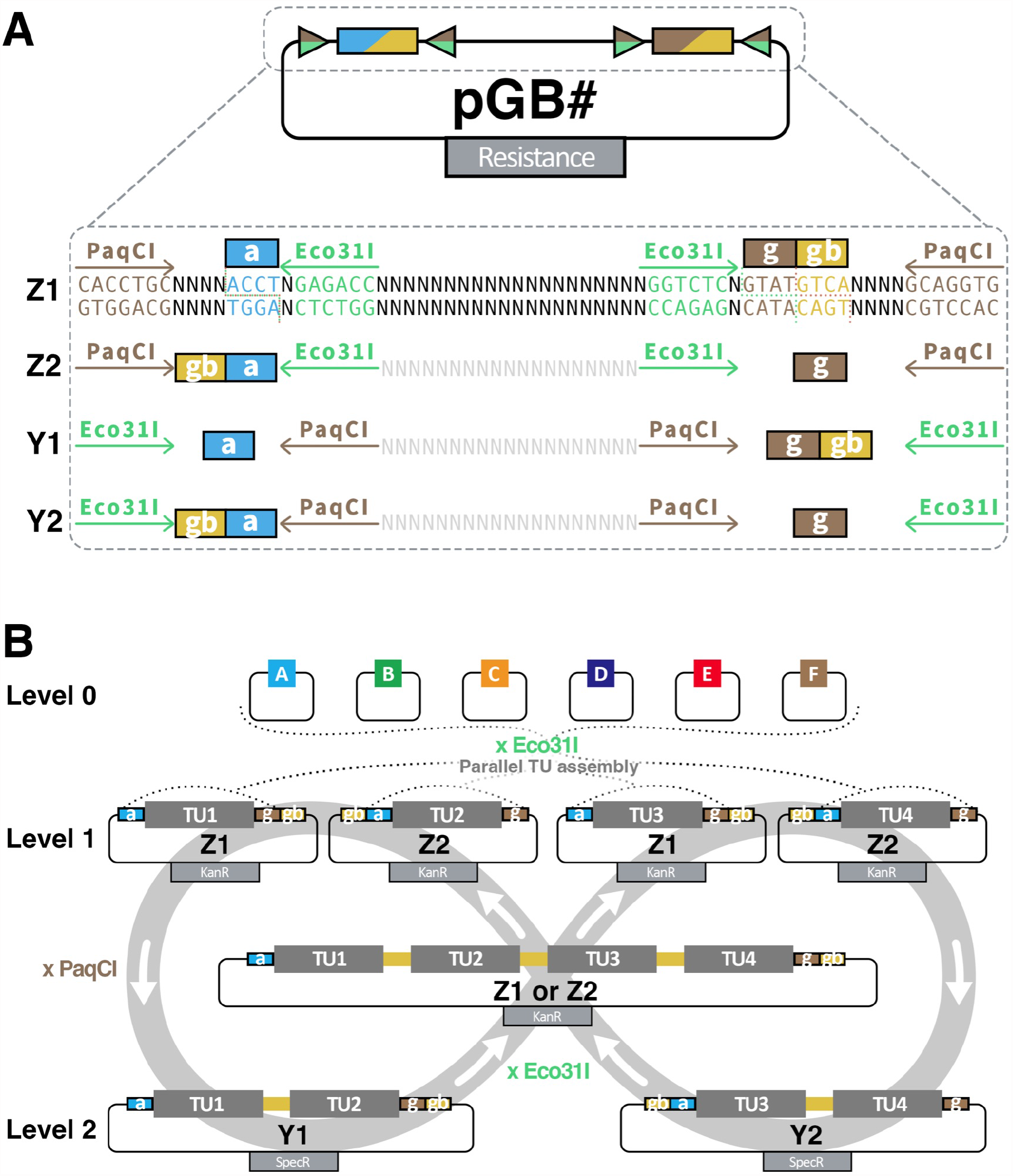
Iterative stacking of TUs with GreenBraid. (A) Key components and cloning sites of the GreenBraid destination plasmids (pGB). The PaqCI/AarI and Eco31I recognition sites are depicted as arrows, and the corresponding overhangs as filled boxes with the overhang in lower letters. (B) Principle of GreenBraid (GB) iterative assembly of multiple TUs on a single vector. Level 0 GG or GG2.0 entry vectors are assembled in parallel by Eco31I GoldenGate reaction in Z1 or Z2 destination vectors to form four different Level 1 TUs (TU1 to TU4). A TU in Z1 and a TU in Z2 can be assembled by PaqCI/AarI GoldenGate reaction in Y1 or Y2 destination vectors to generate a Level 2 multigene assembly. Two of these two-TU Level 2 can be assembled in a Z1 or Z2 destination vector to generate a four-TU construct. These can be further combined with additional TUs according to the same logic.

Four Level 1 vectors (pGB-Zx, Figure 5A, Supplemental Figure S1) carry a kanamycin resistance gene and the spisPink chromoprotein gene as a negative cloning screening marker. spisPink colours the colonies pink; it is released by Eco31I digestion and marks colonies with successfully assembled TUs as white. All Level 1 destination vectors contain the standard “a” (ACCT) and “g” (CGCT) overhang to accommodate a TU, plus the GreenBraid “gb”(GTCA) overhang that allows the concatenation of TUs by switching between Level 1 and Level 2 vectors. The pGB-Z1 and pGB-Z2 Level 1 vectors only differ from the pGB-Z1r and pGB-Z2r variants by the orientation of the “a” and “g” overhangs with respect to the “gb” overhang (Figure 5A, Supplemental Figure S1) and allow to control the respective orientation of the TUs (parallel, converging or diverging).

Four Level 2 vectors (pGB-Yx, Figure 5A, Supplemental Figure S1) carry a spectinomycin resistance gene and the sfGFP chromoprotein gene as a negative cloning screening marker. sfGFP colours the colonies green; it is released by PaqCI/AarI digestion and marks colonies with successfully assembled TUs as white. All Level 2 destination vectors contain the standard “a” (ACCT) and “g” (CGCT) overhangs to accommodate a multi-TU, plus the GreenBraid “gb”(GTCA) overhang, that allows the concatenation of TUs. As for the Level 1 vectors, the pGB-Y1 and pGB-Y2 Level 1 vectors only differ from the pGB-Y1r and pGB-Y2r variants by the orientation of the “a” and “g” overhangs with respect to the “gb” overhang (Figure 5A, Supplemental Figure S1) and allow to control the respective orientation of the TUs.

All Level 1 and Level 2 destination vectors are built on the pCAMBIA backbone [14] and can thus be directly used in plants. It is important to note that TU assemblies from Level 0 can only be done in Level 1 destination vectors (pGB-Z1 or Z2) and not in Level 2 (pGB-Y1 or Y2). GreenBraiding relies on two different type IIs endonucleases. PaqCI/AarI combines two Level 1 vectors in a Level 2. Eco31I combines two Level 2 vectors back to a Level 1 vector. Using the same two endonucleases as in Level 0 and Level 1 assembly ensures that any brick domesticated for these enzymes can be used for multi-TU stacking. We optimised the protocol for a one-pot GoldenGate assembly with PaqCI/AarI in Level 2 reactions for the most successful clones. The optimal protocol uses a higher concentration of T4 ligase and is shorter than a regular Eco31I-based GreenGate assembly of Level 1 (two hours instead of eight hours). The efficiency of GreenBraiding is excellent, with typically 70±26% (mean±sd, n= 33 assemblies) of colonies containing the correctly assembled TUs.

In addition to the above-described core set of Level 1 and Level 2 destination vectors, we generated several auxiliary plasmids (pGBZ1j, Z2j, Y1j, Y2j) containing a dummy 162nt-long insert. These auxiliary plasmids, dubbed “Jokers”, can substitute for TUs during GreenBraiding and enable the assembly of an uneven number of TUs (Supplementary Table S1).

To validate the stacking of TUs by GreenBraiding, we constructed a proof-of-concept plant transformation vector containing four TUs. We aimed to express four fluorescent-tagged fusion proteins expressed from four different promoters in *Arabidopsis thaliana*. Four individual TUs were assembled from Level 0 entry clones (Figure 6A). TU1: a translational fusion between the nucleus-localised Histone 2B (H2B) and mTurquoise2 expressed from the RPS5A promoter active in dividing cells [19]. TU2 is a translational fusion between the plasma membrane-localised aquaporin PIP1;4 [20], and mVenus is expressed from the ubiquitously expressed UBI10 promoter. TU3: a translational fusion between the tonoplast-localized subunit a3 of the vacuolar proton pump (Vha-a3, [21]) and mScarlet expressed from the lateral root promoter pGATA23 [22]. TU4: a translational fusion between Vha-a3 and mScarlet expressed from the xylem pole pericycle promoter pXPP [23]. In all four TUs, a 14 to 37 amino-acid-long Glycine-Serine linker is present between the fluorescent protein and the tagged protein and the same *tHSP18*.*2* terminator was used. These four TUs were combined two-by-two in Level 2 vectors (Figure 6A), and the resulting plasmids were combined again to combine all four TUs in the final Level 1 destination vector. An F module providing FastRed selection [24] was used to assemble TU4, while dummy F modules were used to assemble the other TUs. Thus the final construct only expresses a single copy of the selection marker. GreenBraiding-based stacking of the four TUs was efficient (83%), resulting in a 26kb plasmid verified by sequencing. The plasmid construct was transformed in *Agrobacterium tumefaciens* GV3101, where it remained intact (100% of colonies tested). In plants, ∼5% of primary transformants expressed all four reporters at high levels, where expected (Figure 6D): the plasma membrane marker PIP1;4-mVenus in all tissues, H2B-mTurquoise2 in meristematic cells, the vacuolar marker VHA3-mScarlet in the pericycle cells facing the xylem poles and the cells of the lateral root primordia, but not in the root apical meristem (Figure 6D). This expression was stable across generations. In most T1, the lateral-root-primordia-specific expression of the vacuolar marker was either weak or showed patchy expression (observed in some lateral root primordia but not all).

**Figure 6.**
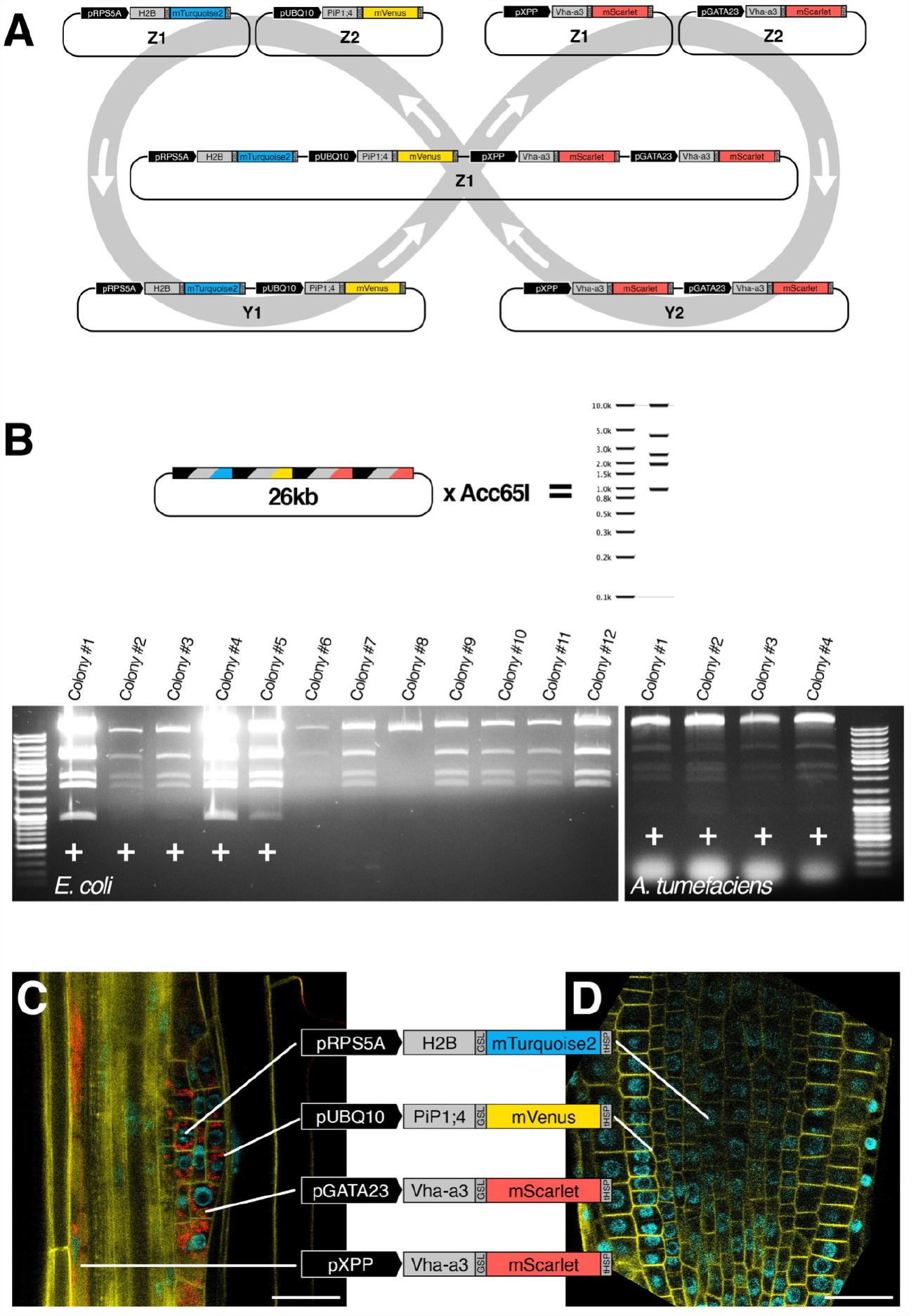
Proof-of-concept assembly and expression of a four TU construct in *Arabidopsis thaliana*. (A) Graphical depiction of the proof-of-concept assembly where a plant transformation vector with four transcriptional units is assembled. Each TU expresses a fluorescent fusion protein that localises to a specific cellular domain. The entry module’s nicknames are noted in the individual transcriptional units. The colour (blue, yellow, or green) corresponds to the colour of emitted light from excited fluorescent proteins (B) Virtual digest of the proof-of-principle plant transformation vector, and two screenings of several *E*.*coli* or *A*.*tumefaciens* colonies for colonies that contain the proof-of-principle plant transformation vector post bacterial transformation. Positive colonies are marked with a (+). (C-D) Sections of the primary root of a transformed *A*.*thaliana* seedling that visualises expression of each transcriptional unit on the proof-of-principle T-DNA. The blue nuclei marker is seen in proliferating cells, the yellow plasma membrane marker marks all cells, and the two red tissue-specific vacuole markers are seen separately in xylem pole pericycle cells or a stage three lateral root primordia.

### A set of pre-assembled Level 1 vectors

Live imaging and targeted manipulation of gene expression frequently relies on fluorescent plasma membrane (PM) and nuclei markers, along with inducible expression systems for transgenes. We have developed a series of pre-constructed, ubiquitously expressed PM markers, nucleus markers, and an LhG4 driver in GreenBraid Level 1 Z vectors (see Supplemental Figure S3). These vectors are readily usable for a Level 2 GreenBraid assembly with any desired construct. The array of fluorescent markers is extensive, encompassing 22 pre-cloned Level 1 constructions that offer great flexibility when devising a cloning strategy. These constructions consist of fluorescent fusion proteins, combining PiP1;4 (PM marker) or H2B (nuclei marker) with one of three fluorescent fusion proteins (mTurquoise2, mVenus, or mScarlet), all driven by the UB10 promoter. The markers were constructed in Z1 and Z2 GreenBraid vectors, and each is available in two versions: one with a plant selection marker (FastRed, a red fluorescent seed selection marker [25]). This comprehensive toolkit empowers researchers with versatile and efficient tools to study various cellular processes.

## Conclusions

GG2.0 overcomes the limitations of the original GreenGate approach. GG2.0 introduces a Universal Entry Generator plasmid (pUEG) that allows one-pot, one-step assembly of Level 0 modules, facilitating easy domestication of internal sites and creating entry clones efficiently. By expanding the repertoire of available overhangs, GG2.0 enables the assembly of Level 1 vectors from up to 12 Level 0 modules, greatly increasing the modularity of GreenGate assemblies. GG2.0 also offers a set of Level 0 entry plasmids for multiplex CRISPR/Cas9 genome editing. This feature allows the generation of T-DNA vectors containing up to 18 gRNAs and a promoter-Cas9-terminator cassette for efficient multiplex genome editing. Additionally, GG2.0 introduces GreenBraiding, a novel method that enables iterative stacking of transcriptional units (TUs) using plant-compatible Level 1 and Level 2 destination vectors. This process facilitates the parallel and efficient two-by-two combination of TUs, offering limitless potential for TU stacking. We also present a set of pre-assembled Level 1 vectors featuring fluorescent plasma membrane (PM) and nuclei markers, and for LhG4-driven inducible gene expression. These vectors are readily usable for Level 2 GreenBraid assembly with any desired construct, providing a comprehensive toolkit for researchers to study various cellular processes. GreenGate 2.0 is significantly improved over the original GreenGate, while being backwards compatible. It significantly broadens the capabilities and applications of the GreenGate system, enabling precise and efficient genetic engineering in plant research.

## Supporting information

- **Figure S1**. Schematic overview of all GB destination plasmids and their name
- **Figure S2**. Schematic overview of all Level 0 plasmids for multiplex expression of gRNAs for CRISPR/Cas9 genome editing and their name.
- **Figure S3**. Schematic overview of all filled GB destination plasmids and their name
- **Table S1**. The GreenGate 2.0 vector set
- **Table S2**. Oligonucleotides used in this study
- **Dataset S1**. Sequences of all GreenGate 2.0 vectors (gene bank format)
- **Supplemental Protocol 1**. Creating entry GG clones and Domestication using the pUEG
- **Supplemental Protocol 2**. GreenGate assembly of Level 0 modules into a Level 1 vector
- Supplemental Protocol 3. GreenBraiding

## Data Availability Statement

All relevant data are within the paper and its Supporting Information files.

## Acknowledgements

We thank Elif Gediz Kocaoglan for the chromoproteins clones (spisPink & sfGFP), to Uriel Urquiza & Miguel Miñambres Martín for the GoldenBraid vectors and Robertas Ursache for their gRNA expression vectors. Our gratitude goes to Celine Geiger, Freddy Can Igiebor, Raffaela Nauke, Charlotte Neumann, Cathrin Rube, Gülce Tekin and Caja-Marie Winzer students of the Molecular and Applied Plant Science Master program at Heidelberg University, for their help in generating the pre-assembled Level 1 plasmids with plasma membrane markers.

## Author Contributions

Conceptualisation: Marcel Piepers, Alexis Maizel

Funding acquisition: Tomas Tessi, Alexis Maizel

Investigation: Marcel Piepers, Katarina Erbstein, Jazmin Reyes-Hernandez, Changzheng Song, Tomas Tessi, Vesta Petrasiunaite, Naja Faerber, Kathrin Distel

Methodology: Marcel Piepers, Katarina Erbstein, Alexis Maizel

Resources: Marcel Piepers, Katarina Erbstein, Jazmin Reyes-Hernandez, Changzheng Song, Tomas Tessi, Naja Faerber, Kathrin Distel, Alexis Maizel

Supervision: Alexis Maizel

Writing – original draft: Marcel Piepers, Alexis Maizel

Writing – review & editing: Marcel Piepers, Katarina Erbstein, Alexis Maizel

## Footnotes

1 Submission of the plasmids set to AddGene has been initiated. The accession will be provided when completed.

## References

1. Patron NJ, Orzaez D, Marillonnet S, Warzecha H, Matthewman C, Youles M, et al. Standards for plant synthetic biology: a common syntax for exchange of DNA parts. New Phytol. 2015;208: 13–19. doi:10.1111/nph.13532

2. Kirchmaier S, Lust K, Wittbrodt J. Golden GATEway cloning--a combinatorial approach to generate fusion and recombination constructs. PLoS One. 2013;8: e76117. doi:10.1371/journal.pone.0076117

3. Gibson DG, Young L, Chuang R-Y, Venter JC, Hutchison CA 3rd, Smith HO. Enzymatic assembly of DNA molecules up to several hundred kilobases. Nat Methods. 2009;6: 343–345. doi:10.1038/nmeth.1318

4. Sleight SC, Bartley BA, Lieviant JA, Sauro HM. In-Fusion BioBrick assembly and re-engineering. Nucleic Acids Res. 2010;38: 2624–2636. doi:10.1093/nar/gkq179

5. Bird JE, Marles-Wright J, Giachino A. A User’s Guide to Golden Gate Cloning Methods and Standards. ACS Synth Biol. 2022;11: 3551–3563. doi:10.1021/acssynbio.2c00355

6. Lampropoulos A, Sutikovic Z, Wenzl C, Maegele I, Lohmann JU, Forner J. GreenGate---a novel, versatile, and efficient cloning system for plant transgenesis. PLoS One. 2013;8: e83043. doi:10.1371/journal.pone.0083043

7. Sarrion-Perdigones A, Falconi EE, Zandalinas SI, Juárez P, Fernández-del-Carmen A, Granell A, et al. GoldenBraid: an iterative cloning system for standardized assembly of reusable genetic modules. PLoS One. 2011;6: e21622. doi:10.1371/journal.pone.0021622

8. Weber E, Engler C, Gruetzner R, Werner S, Marillonnet S. A modular cloning system for standardized assembly of multigene constructs. PLoS One. 2011;6: e16765. doi:10.1371/journal.pone.0016765

9. Engler C, Youles M, Gruetzner R, Ehnert T-M, Werner S, Jones JDG, et al. A golden gate modular cloning toolbox for plants. ACS Synth Biol. 2014;3: 839–843. doi:10.1021/sb4001504

10. Andreou AI, Nakayama N. Mobius Assembly: A versatile Golden-Gate framework towards universal DNA assembly. PLoS One. 2018;13: e0189892. doi:10.1371/journal.pone.0189892

11. Andreou AI, Nirkko J, Ochoa-Villarreal M, Nakayama N. Mobius Assembly for Plant Systems highlights promoter-terminator interaction in gene regulation. bioRxiv. bioRxiv; 2021. p. 2021.03.31.437819. doi:10.1101/2021.03.31.437819

12. Chamness JC, Kumar J, Cruz AJ, Rhuby E, Holum MJ, Cody JP, et al. An extensible vector toolkit and parts library for advanced engineering of plant genomes. Plant Genome. 2023; e20312. doi:10.1002/tpg2.20312

13. Pollak B, Matute T, Nuñez I, Cerda A, Lopez C, Vargas V, et al. Universal loop assembly: open, efficient and cross-kingdom DNA fabrication. Synth Biol. 2020;5: ysaa001. doi:10.1093/synbio/ysaa001

14. Sarrion-Perdigones A, Vazquez-Vilar M, Palací J, Castelijns B, Forment J, Ziarsolo P, et al. GoldenBraid 2.0: a comprehensive DNA assembly framework for plant synthetic biology. Plant Physiol. 2013;162: 1618–1631. doi:10.1104/pp.113.217661

15. Green MR, Sambrook J. Preparation of Plasmid DNA by Alkaline Lysis with Sodium Dodecyl Sulfate: Minipreps. Cold Spring Harb Protoc. 2016;2016. doi:10.1101/pdb.prot093344

16. Ursache R, Fujita S, Dénervaud Tendon V, Geldner N. Combined fluorescent seed selection and multiplex CRISPR/Cas9 assembly for fast generation of multiple Arabidopsis mutants. Plant Methods. 2021. p. 2021.05.20.444986. doi:10.1186/s13007-021-00811-9

17. Clough SJ, Bent AF. Floral dip: a simplified method for Agrobacterium-mediated transformation of Arabidopsis thaliana. Plant J. 1998;16: 735–743. doi:10.1046/j.1365-313x.1998.00343.x

18. Stuttmann J, Barthel K, Martin P, Ordon J, Erickson JL, Herr R, et al. Highly efficient multiplex editing: one-shot generation of 8× Nicotiana benthamiana and 12× Arabidopsis mutants. Plant J. 2021;106: 8–22. doi:10.1111/tpj.15197

19. Tsutsui H, Higashiyama T. pKAMA-ITACHI vectors for highly efficient CRISPR/Cas9-mediated gene knockout in Arabidopsis thaliana. Plant Cell Physiol. 2016; cw191. doi:10.1093/pcp/pcw191

20. Geldner N, Dénervaud-Tendon V, Hyman DL, Mayer U, Stierhof Y-D, Chory J. Rapid, combinatorial analysis of membrane compartments in intact plants with a multicolor marker set. Plant J. 2009;59: 169–178. doi:10.1111/j.1365-313X.2009.03851.x

21. Dettmer J, Hong-Hermesdorf A, Stierhof Y-D, Schumacher K. Vacuolar H+-ATPase activity is required for endocytic and secretory trafficking in Arabidopsis. Plant Cell. 2006;18: 715–730. doi:10.1105/tpc.105.037978

22. De Rybel B, Vassileva V, Parizot B, Demeulenaere M, Grunewald W, Audenaert D, et al. A novel aux/IAA28 signaling cascade activates GATA23-dependent specification of lateral root founder cell identity. Curr Biol. 2010;20: 1697–1706. doi:10.1016/j.cub.2010.09.007

23. Andersen TG, Naseer S, Ursache R, Wybouw B, Smet W, De Rybel B, et al. Diffusible repression of cytokinin signalling produces endodermal symmetry and passage cells. Nature. 2018;555: 529–533. doi:10.1038/nature25976

24. Vilches Barro A, Stöckle D, Thellmann M, Ruiz-Duarte P, Bald L, Louveaux M, et al. Cytoskeleton Dynamics Are Necessary for Early Events of Lateral Root Initiation in Arabidopsis. Curr Biol. 2019;29: 2443–2454.e5. doi:10.1016/j.cub.2019.06.039

25. Shimada TL, Shimada T, Hara-Nishimura I. A rapid and non-destructive screenable marker, FAST, for identifying transformed seeds of Arabidopsis thaliana. Plant J. 2010;61: 519–528. doi:10.1111/j.1365-313X.2009.04060.x

